# Comparative primate transcriptomics identifies a ZNF90–OVOL2 regulatory axis shaping human neural progenitor cell dynamics

**DOI:** 10.64898/2026.07.27.740929

**Authors:** Cesar Mateo Bastidas-Betancourt, Maria Isabella Negretti-Ingravallo, İrfan Burak Göloğlu, Dilnaz Begenova, Mireia S. Puig-Segui, André Mendes Costa, Mariya Reshetar, Sabrina Heide, Martin N. Ivanov, Lora V. Veleva, Tsvetomir E. Kachovski, Emil G. Kovachev, Anton B. Tonchev, Michael Heide

**Author notes:** first author (Cesar Mateo Bastidas-Betancourt).

## Abstract

Understanding the behavior of human neural progenitor cells (NPCs) requires comparative characterization of their transcriptomic landscape, particularly through comparisons with closely related primate species. The rhesus macaque represents a key non-hominoid outgroup for such analyses, and several representative transcriptomic datasets are now available. Here, we compare genes enriched in human NPCs with those enriched in rhesus macaque NPCs and found that human NPC-enriched genes are associated with gene programs supporting radial glial identity and proliferative capacity, as well as functions related to sister chromatid segregation. This analysis identified two zinc-finger transcription factors for functional investigation: the ape-specific ZNF90 and the highly conserved OVOL2. By analysing their genome-wide binding, transcriptional output, and cellular effects in cerebral organoids, we uncovered a previously uncharacterized regulatory axis with divergent but partially overlapping effects that converge on apical progenitor (AP) maintenance. Together, our findings support a model in which lineage-specific / evolutionarily young transcription factor become integrated into a conserved developmental gene regulatory network, generating novel NPC dynamics during primate corticogenesis.

## Introduction

The expanded neocortex is a defining feature of the human brain^1–4^. At the cellular level, this evolutionary expansion has been linked to changes in both the duration and tempo of neurogenesis^5^ and to an increased size and proliferative capacity of the neural progenitor cell (NPC) pool^6,7^. In humans, the neurogenic period is prolonged compared with that of other primates, allowing cortical neurons to be generated over an extended period of fetal development^8^. In addition, the human fetal neocortex contains an expanded population of NPCs, which forms an essential basis for increased neuronal output^6^. Such evolutionary changes are thought to arise, at least in part, from species-specific differences in NPC behavior. During early corticogenesis, NPCs initially line the ventricular zone (VZ) as neuroepithelial cells (NECs), which divide symmetrically before undergoing the neuroepithelial transition (NET) to form apical radial glia (aRGs)^9,10^. Together, NECs and aRGs constitute the apical progenitor (AP) pool. During corticogenesis, APs either generate neurons directly or produce basal progenitors (BPs), which delaminate from the VZ and populate the subventricular zone (SVZ), where they generate neurons indirectly^11,12^. BPs comprise basal intermediate progenitors (bIPs) and basal radial glia (bRG), the latter of which are particularly abundant in gyrencephalic mammals, including primates, and have been strongly associated with neocortical expansion^13–15^.

The gene regulatory networks (GRNs) governing NPC behavior remain incompletely understood. Comparative transcriptomic studies have identified numerous developmental regulators and several lineage-specific genes that likely contributed to primate brain evolution. However, considerably less is known about how newly evolved regulatory factors become incorporated into pre-existing developmental GRNs that have been conserved across mammals. Because NPC behavior is orchestrated by deeply conserved transcriptional programs, evolutionary innovation is likely to depend not only on the emergence of novel regulators but also on their functional integration into these pre-existing regulatory programs. Identifying how evolutionarily conserved and lineage-specific transcription factors cooperate within shared GRNs therefore represents an important step toward understanding the molecular basis of primate brain evolution. One powerful approach to identify these networks and their upstream regulators is transcriptomic profiling of human fetal brain tissue with a focus on NPC populations. Such studies have uncovered key regulators of NPC proliferation, differentiation, and lineage progression, including several human-specific genes, such as *ARHGAP11B*, and members of the *NOTCH2NL* and *NBPF* gene families, that have likely contributed to the evolutionary expansion of the human neocortex^16–21^. However, understanding how these GRNs evolved requires comparison with closely related primates possessing smaller and less folded brains. In this context, the rhesus macaque (*Macaca mulatta*) represents a particularly informative comparative model. As an Old-World monkey (Catarrhini), it occupies a key phylogenetic position outside the hominoid lineage while sharing many fundamental aspects of cortical development with humans. Comparative studies of rhesus corticogenesis have therefore been instrumental in reconstructing conserved features of NPC behavior and developmental timing among catarrhine primates^4,13,22,23^. Despite its importance, comparatively few studies have functionally examined the regulatory mechanisms controlling NPC behavior across humans and rhesus macaques.

Transcription factors are central organizers of GRNs because they coordinate large transcriptional programs controlling cell identity and developmental transitions. Consequently, evolutionary changes in transcription factor repertoires may not only generate novel regulatory activities but also reshape existing GRNs through interactions with conserved developmental regulators. One of the largest transcription factor families in the animal kingdom is the zinc finger protein (ZNF) family, which has undergone remarkable expansion during mammalian evolution and particularly throughout primate evolution^24–27^. ZNFs are characterized by C2H2 zinc finger domains that mediate sequence-specific DNA binding^28,29^. This family includes numerous well-established regulators of neural development, including GLI3, FEZF1, FEZF2, and ZEB1. Moreover, several ZNF family members, including ZNF238, ZNF519, and ZNF568, have been implicated in cortical malformations^30,31,32^, highlighting the importance of this transcription factor family for normal cortical development and suggesting that additional ZNFs may contribute to the regulation of NPC behavior.

Until recently, experimentally testing evolutionary differences in GRNs between human and rhesus macaque NPCs has been technically challenging. Cerebral organoids (COs) provide a powerful experimental platform to address these questions, as they recapitulate key aspects of neural NPC proliferation, migration, and neuronal differentiation^33,34^, while enabling functional genetic experiments that are not feasible in fetal primate tissue^33,35,36^. We recently established a unified protocol to generate COs from human, chimpanzee, rhesus macaque, and common marmoset induced pluripotent stem cells (iPSCs)^37^. In that study and in subsequent work^38,39^, we demonstrated that microinjection and electroporation of primate COs provide a robust platform for functional interrogation of candidate genes during early primate corticogenesis.

Here, we sought to identify transcriptional regulators contributing to species differences in early primate corticogenesis by re-analyzing published transcriptomic datasets from human and rhesus macaque NPCs. This analysis identified the ape-specific KRAB zinc finger protein ZNF90 and the conserved transcription factor OVOL2 as candidate regulators of NPC biology. We characterized their expression patterns, genomic binding profiles, and individual as well as combined transcriptional and cellular functions. Together, our findings indicate that an evolutionarily young transcription factor can converge with an ancient developmental regulator on shared gene regulatory networks controlling NPC behavior, providing a mechanistic framework for how lineage-specific regulatory innovations become integrated into conserved developmental programs during primate corticogenesis.

## Results

### Comparative transcriptomic analysis identifies human NPC-specific gene programs linked to (apical) radial glial maintenance and chromatid segregation

In order to identify genes preferentially expressed in NPCs and more highly expressed in human than in rhesus macaque NPCs, we first defined NPC-enriched genes in human by analyzing four published bulk RNA-seq datasets of fetal human neocortical tissue^16,18,40^ and of NPCs differentiated from induced pluripotent stem cells^41^, similar to Florio et al. 2018^17^. Specifically, these comprise: (i) the datasets of microdissected germinal zones (VZ, iSVZ and oSVZ) and cortical plate (CP) from Fietz et al. 2012^40^, which we analyzed to identify genes more highly expressed in germinal zones than in the CP; (ii) the dataset of FACS-enriched radial glia cells (aRG and bRG) and neurons from Florio et al. 2015^16^, which we analyzed to identify genes more highly expressed in radial glia cells than in neurons; (iii) the dataset of FACS-enriched radial glia cells (aRG and bRG), intermediate progenitors (IPs), and neurons from Johnson et al. 2015^18^, which we analyzed to identify genes more highly expressed in radial glia cells than in IPs and neurons; and (iv) the dataset of NPCs differentiated from induced pluripotent stem cells from the ENCODE Project Consortium ^41^, which we used as a baseline for genes expressed in human NPCs (Fig. 1A). By integrating these datasets and performing a differential gene expression (DGE) analysis, we identified a set of 2,265 genes upregulated in NPCs (Fig. 1B, Table S2). This gene set strongly overlaps with the gene set by Florio et al. 2018 (Fig. 1C), validating our approach, and is enriched for functional categories related to DNA replication and cell cycle processes (Fig. 1D), consistent with the proliferative state of NPCs. Notably, these genes show significantly greater proximity (<100 kb) to annotated human-accelerated regions^42^ (HARs) compared to other genes in the dataset (Fig. 1E), suggesting an enrichment of HAR-associated regulatory landscapes among NPC-expressed genes.

**Figure 1.**
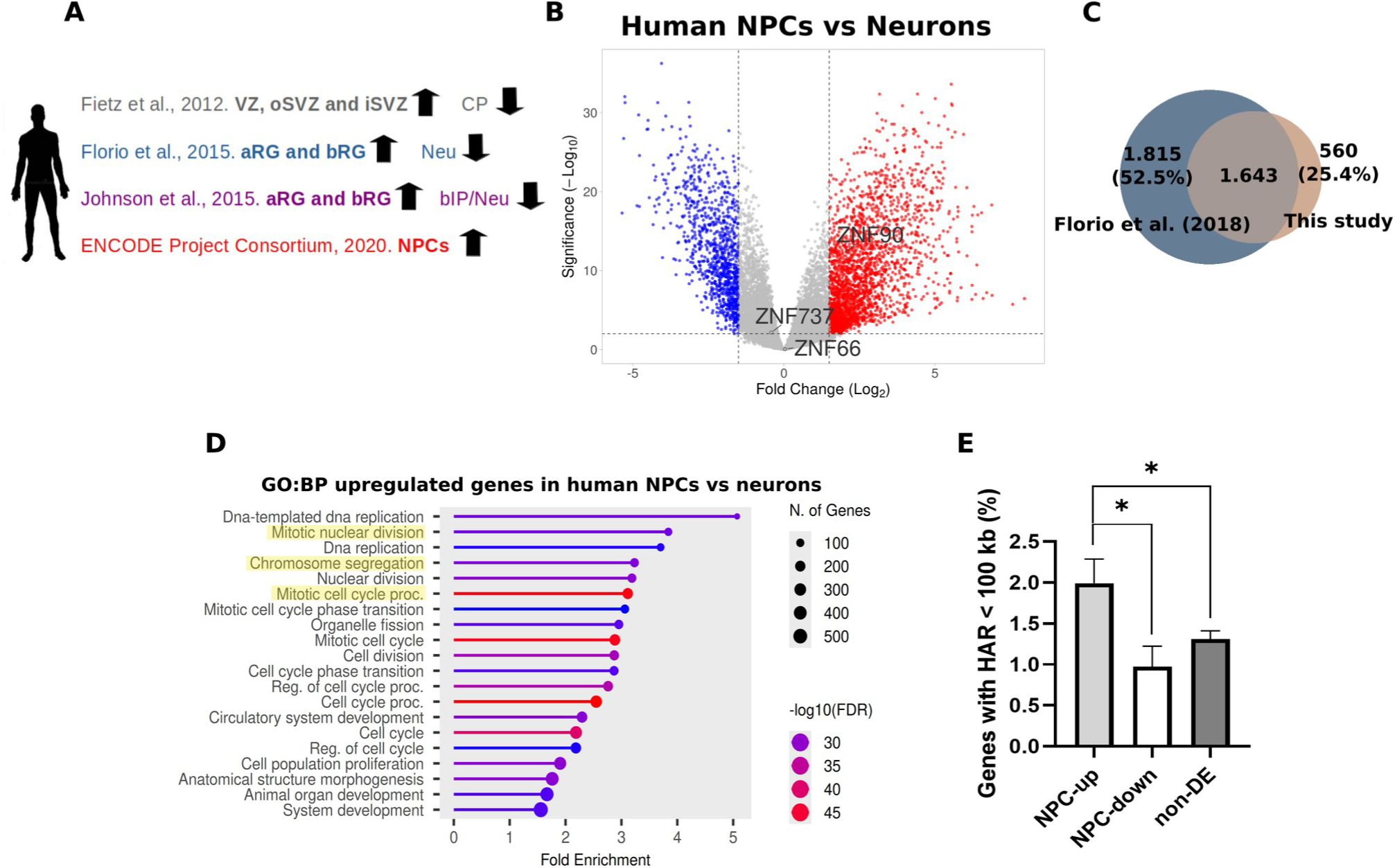
Comparative transcriptomic analyses define a human neural progenitor gene signature enriched for cell cycle-associated functions and human-accelerated regulatory regions. (A) Transcriptome datasets of fetal human neocortical tissue and of human NPCs differentiated from induced pluripotent stem cells, reanalyzed for genes more highly expressed in NPCs/germinal zones (upward arrows) than in neurons/cortical plate (downward arrows). (B) Volcano plot showing log2 fold change (log2FC; x-axis) and statistical significance (-log10 FDR; y-axis) for genes identified in the re-analysis, highlighting ZNF90 and its two closest paralogues, ZNF737 and ZNF66. (C) Venn diagram showing the overlap between NPC-enriched genes reported by Florio et al. (2018)^17^ and those identified in this study. (D) GO biological process (GO:BP) enrichment analysis of genes overexpressed in human NPCs compared with neurons (n = 2,265 genes). (E) Percentage of genes located close to a HAR (<100 kb) among genes upregulated in NPCs (NPC-up), downregulated in NPCs (NPC-down), and non-differentially expressed genes (non-DE). Statistical significance was assessed using Fisher’s exact test. Asterisks indicate significant differences in HAR abundance (P < 0.05). Exact P-values: NPC-up versus NPC-down, P = 0.016; NPC-up versus non-DE, P = 0.015.

In a next step, we asked whether comparable bulk RNA-seq datasets are available for rhesus macaque NPCs. We identified three such datasets: (i) the dataset of microdissected germinal zone and CP from Luo et al. 2021^43^, which we analyzed to identify genes more highly expressed in the germinal zone than in the CP; (ii) the dataset of NPCs and rosette neural stem cells generated from rhesus macaque embryonic stem cells from Zhao et al. 2014^44^, which we used as a baseline for genes expressed in rhesus macaque NPCs; and (iii) the dataset of isolated and cultured NPCs from prenatal rhesus macaque telencephalon from Micali et al. 2023^45^, which we used as a baseline for genes expressed in rhesus macaque NPCs (Fig. 2A). By integrating these datasets and performing a cross-species DGE analysis based on liftover mapping of rhesus macaque genes to the human genome annotation^46^, we identified a set of 1.404 genes showing human NPC enrichment, defined as genes for which the NPC-versus-neuron expression difference was greater in human than in rhesus macaque. These genes were enriched for functions related to chromatid segregation, cell division, and vascular development (Fig. 2B). Rather than reflecting bona fide endothelial contamination, the vascular development signature likely represents a shared apical polarity and mechanosensory gene program characteristic of radial glia, associated with maintenance of ventricular zone identity and proliferative capacity. Consistent with this interpretation, genes within this category include markers associated with aRG organization (*CLDN5, PECAM1, EMCN, HEG1, ROBO4, TIE1, TEK, KDR, FLT4, SOX17, SOX18, GATA2*), NPC maintenance signaling pathways (*HES1, DLL4, NOTCH4, NRARP*), mechanotransduction and adhesion-associated regulators (*YAP1, ITGB1BP1, S1PR1, PTK2B*), and growth factor–mediated niche signaling (*PDGFB, VEGFB, PGF, FGF1*) (Table S3). Moreover, weighted gene co-expression network analysis (WGCNA) performed on the same datasets revealed human NPC-associated modules enriched for genes involved in DNA replication and cell cycle regulation (Fig. 2C), with canonical cell-cycle genes such as *CDKN1B, RMI2, MCM4*, and *MCMBP* among the most highly connected nodes, indicating a hub-like behavior within the human NPC network (Fig. S1A). Taken together, these data indicate that, compared to rhesus macaque NPCs, human NPCs exhibit enhanced engagement of gene networks associated with chromatid segregation fidelity, as well as reinforced programs supporting maintenance of radial glial identity and proliferative capacity.

**Figure 2.**
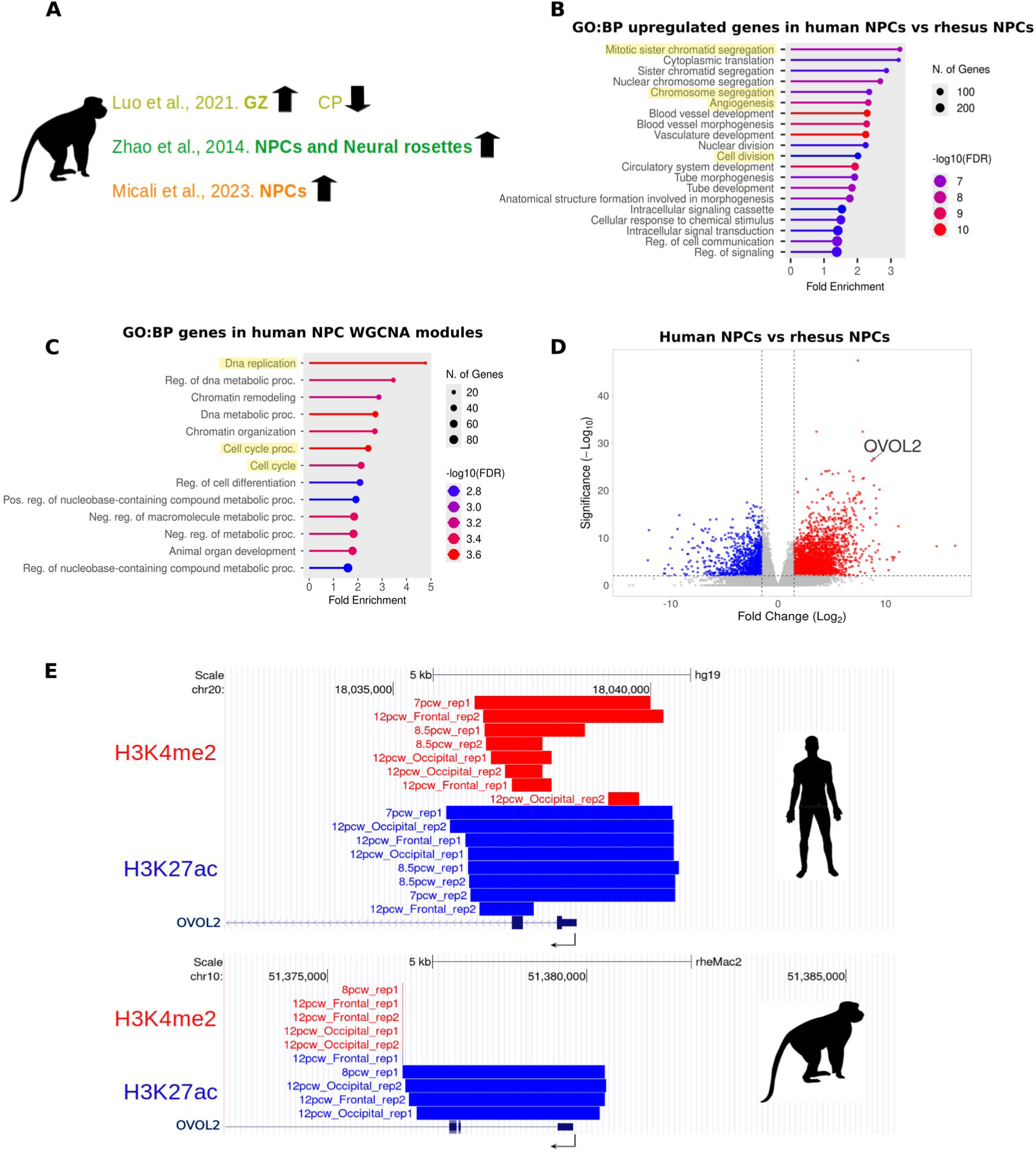
Cross-species transcriptomic analyses identify OVOL2 and ZNF90 as candidate regulators preferentially expressed in human neural progenitor cells. (A) Transcriptome datasets of fetal rhesus macaque neocortical tissue and of rhesus macaque NPCs differentiated from embryonic stem cells and isolated and cultured from prenatal telencephalon, reanalyzed for genes more highly expressed in NPCs/germinal zones (upward arrows) than in neurons/cortical plate (downward arrows). (B) GO:BP enrichment analysis of genes showing stronger NPC enrichment in human than in rhesus macaque, based on the cross-species comparison of NPC-versus-neuron contrasts (n = 1,404 genes). (C) GO:BP enrichment analysis of genes contained in the two WGCNA modules enriched exclusively in human NPC samples. (D) Volcano plot showing log2 fold change (log2FC; x-axis) and statistical significance (-log10 FDR; y-axis) for genes identified in the cross-species re-analysis, highlighting OVOL2 among human NPC-enriched genes. (E) UCSC genome tracks of human (top, hg19) and rhesus macaque (bottom, rheMac2) PCW 5-12 cortical samples from Reilly et al. (2015)^53^, showing H3K4me2 (red) and H3K27ac (blue) ChIP-seq peaks around the core promoter of the corresponding OVOL2 orthologue.

### Identification and validation of conserved and great-ape-specific ZNF candidates, OVOL2 and ZNF90, in human NPCs

Having established a dataset of genes preferentially expressed in NPCs and more highly expressed in human than in rhesus macaque NPCs, we next sought to identify Zinc finger transcription factor (ZNFs) as primary candidates for functional analysis. First, we screened this dataset for ZNFs exhibiting the largest fold changes and most significant p-values. Among these candidates, we prioritized OVOL2 because it combined robust upregulation in human NPCs with known functions in cell-cycle regulation, EMT, and epithelial progenitor state control (Fig. 2D). OVOL2 is a highly conserved metazoan zinc finger transcription factor previously implicated in cell cycle regulation in cancer contexts^47,48^ and in the stabilization of epithelial identity through suppression of epithelial-to-mesenchymal transition (EMT)-like programs^49^. These functions align closely with the key features identified in human NPCs in this study, namely enhanced regulation of chromatid segregation, maintenance of NPC identity, and sustained proliferative capacity. Together, this makes OVOL2 a prime candidate for functional analysis. Second, recent studies have highlighted that clade-specific genes can play important roles in primate brain evolution^50,51^. In contrast to genes showing differential expression, such genes differ in their presence or absence in the genome. We therefore screened our human NPC-enriched gene set for ZNFs present in apes but absent in rhesus macaque. This analysis identified two candidates: the great ape-specific gene *ZNF90* (Fig. S1B) and the human-specific gene *ZNF492*, both previously reported in Florio et al. 2018^17^. We chose to focus on *ZNF90*, as *ZNF492*—due to its more recent evolutionary emergence—is more likely to contribute to differences among great apes rather than between humans and rhesus macaques. In contrast to OVOL2, the functional role of ZNF90 remains largely unexplored, particularly in the context of NPC biology.

To validate the preferential expression of *ZNF90* in NPCs, we analyzed the expression of its closest paralogs, *ZNF737* and *ZNF66* (Fig. S1C,D), in our human NPC dataset and found no significant upregulation in human NPCs relative to neurons, supporting a specific expression pattern of *ZNF90* in the NPC population. Moreover, we detected stable expression of *ZNF90* at early neurodevelopmental stages in human COs (Fig. S1E), consistent with sustained early developmental expression dynamics. Finally, we assessed the expression of *ZNF90* and *OVOL2* in the human immortalized RenCX NPC cell line^52^ and in iPSC-derived human neurons using quantitative real-time PCR (qRT-PCR), confirming significant upregulation of both genes in NPCs, consistent with our RNA-seq reanalysis (Fig. S2A).

To validate the preferential expression of *OVOL2* in NPCs and its increased expression relative to rhesus macaque NPCs, we first analyzed the *OVOL2* core promoter across primate species by multiple sequence alignment and identified a differential expansion of a GC-rich trinucleotide microsatellite (TCCₙ), with 12 repeats in human, 4 in chimpanzee and gorilla, and none in rhesus macaque (Fig. S2B). To further evaluate the regulatory context of *OVOL2* in human and rhesus macaque, we reanalyzed H3K4me2 (accessible chromatin) and H3K27ac (active cis-regulatory elements) ChIP-seq datasets from human and rhesus macaque post conceptional week (PCW) 5–12 cortical tissue^53^. This analysis revealed increased H3K27ac signal at the *OVOL2* locus in human samples (Fig. 2E; Fig. S3). In addition, H3K27ac replicated peaks from human H1-derived NPCs and day-30 COs were consistently detected at the *OVOL2* locus (UCSC track: https://www.genome.ucsc.edu/s/cbastid/H3K27ac%20in%20human%20NPCs%20and%20d30%20organoids), supporting active regulatory engagement during early human neurodevelopment but not in rhesus macaque.

We next examined *OVOL2* expression in a developmental context using human COs and isochronic rhesus macaque COs. *OVOL2* was consistently upregulated in human COs from day 3 after Matrigel embedding (e+3) through day 30 of development (Fig. S4A), a stage at which neurogenesis is established^37,54^. This developmental window corresponds approximately to PCW 13 in human and E70 in rhesus macaque brain development^55^. Notably, *OVOL2* exhibited an expression pattern opposite to that of *ZEB2* (Fig. S4A), a known inhibitory target of OVOL2^56^ and previously implicated in human brain evolution^57^. Comparison of pseudobulk expression with a New World monkey species, the common marmoset^54^, revealed that 30-day-old human COs consistently more highly express *OVOL2* across AP clusters than marmoset COs (Fig. S4B). Finally, *in situ* hybridization analysis of fetal human neocortex (PCW 14) showed strong *OVOL2* expression in the VZ and SVZ, with detectable expression also in the cortical plate (CP) and the marginal zone (MZ) (Fig. S4C). Together, these analyses further support OVOL2 and ZNF90 as robust candidates for functional investigation.

### Genome-wide binding profiles of OVOL2 and ZNF90 reveal promoter-enriched and partially shared binding linked to neurodevelopmental networks

To define the regulatory programs controlled by OVOL2 and ZNF90 in NPCs, we performed CUT&RUN sequencing using N-terminal HA-tagged constructs, analyzing OVOL2 in rhesus macaque COs and ZNF90 in human COs. Genome-wide binding analysis, exemplified by the *DLX1* locus in human for ZNF90 and by the *ZEB1* locus in rhesus macaque for OVOL2 (Fig. S5A–B), revealed enriched binding of both transcription factors near transcriptional start sites (TSSs) and a motif enrichment for the ZNF90-bound regions of stemness TFs such as SOX2/3 and ZIC3 (Figure S5C-D, Table S4). Moreover, both proteins predominantly associate with non-repetitive, gene-proximal DNA regions, including promoters (ZNF90: 7.05%; OVOL2: 8.64%), exons (4.88%; 11.04%), and transcription termination sites (TTS; 5.17%; 2.24%), while being relatively depleted from intronic regions (Fig. 3A,B, Table S5). In addition, both factors exhibited binding to repetitive elements, including human endogenous retroviruses (HERVs), Alu elements, and THE1/LTR sequences (Fig. 3C, Table S6), consistent with previously described DNA-binding properties of zinc finger proteins^58^. Notably, OVOL2 exhibited a substantially larger number of binding sites than ZNF90 (91.983 vs. 6.599 peaks).

**Figure 3.**
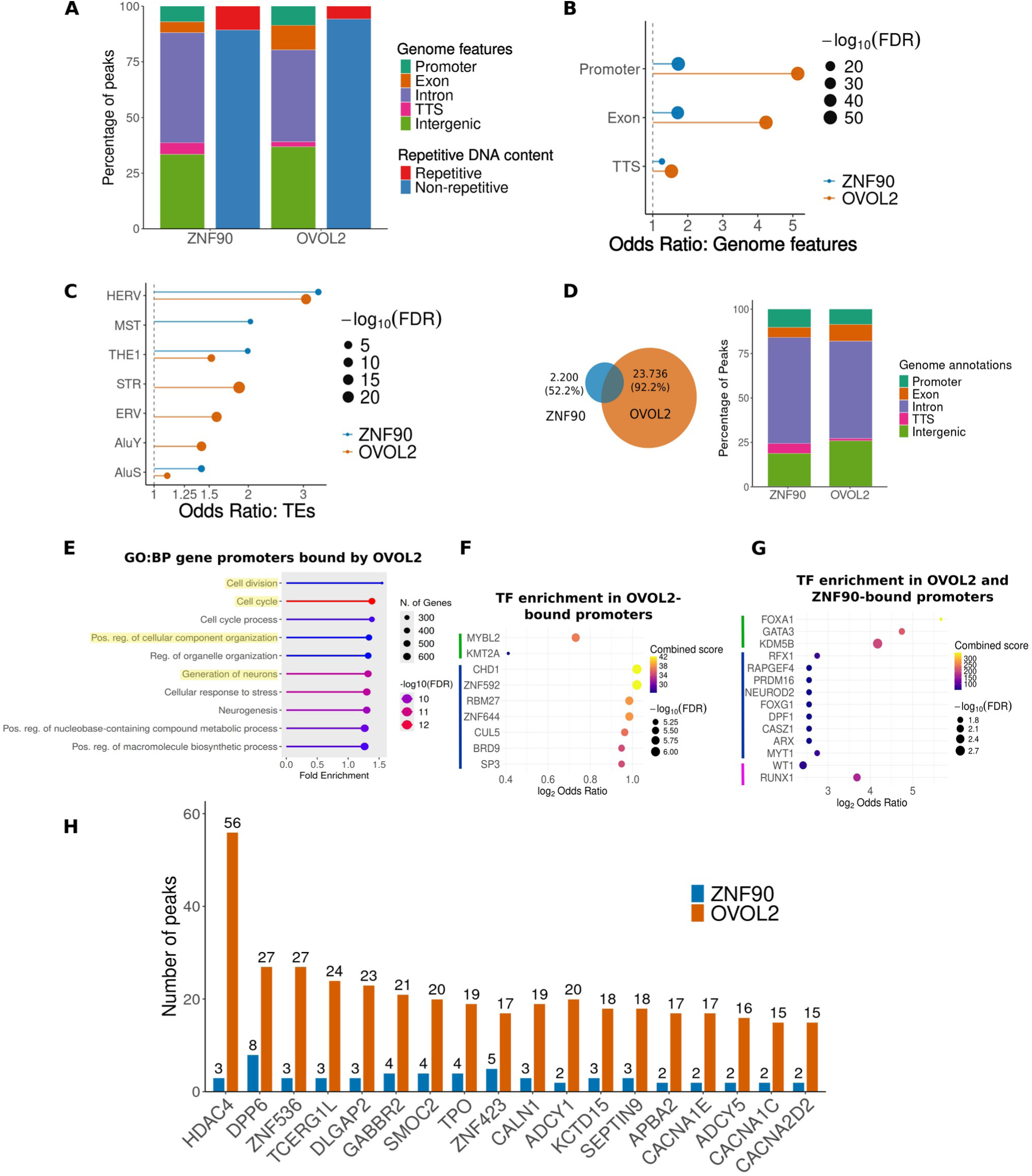
OVOL2 and ZNF90 converge on promoter-associated regulatory networks while retaining distinct target specificities. (A) Mapping percentages for the 6,599 ZNF90 and 91,983 OVOL2 peaks according to their genomic feature annotation (left) and genomic repetitiveness (right). (B and C) Lollipop plots showing the odds ratio enrichment of gene-associated features (B) and TEs (C) for ZNF90 (blue) and OVOL2 (orange), with corresponding statistical significance indicated by circle size. Odds ratios were calculated using Fisher’s exact test, followed by Benjamini–Hochberg correction; circle size indicates adjusted statistical significance. (D) Overlap (n = 2,011 genes) between the 4,211 and 25,747 genes associated with all annotated ZNF90 and OVOL2 peaks, respectively (left), and genomic features associated with these genes for ZNF90 and OVOL2 (right). (E) GO:BP enrichment of genes whose promoters are bound by OVOL2 (n = 7,762). (F and G) Lollipop plots showing odds ratio enrichment and statistical significance for neurodevelopmental TF signatures for OVOL2-bound promoters (F) and the 100 commonly bound gene promoters (G), according to Enrichr’s Rummagene TFs (green line), TF-Gene Co-occurrence Submissions (blue line), and Protein–Protein Interactions (PPIs) (magenta line) TF databases. Odds ratios were calculated using Fisher’s exact test, followed by Benjamini–Hochberg correction; circle size indicates adjusted statistical significance. (H) Number of peaks mapping around genes with more than one peak bound by both ZNF90 (blue) and OVOL2 (orange).

Comparison of target sets revealed substantial overlap, with 47.8% of the 4,211 ZNF90 target genes shared with OVOL2, whereas only 7.8% of the 25,747 OVOL2 targets genes overlapped with ZNF90 (Fig. 3D). We first analyzed factor-specific peaks. Promoter-associated OVOL2-linked targets (7.767) were enriched for functional categories related to cell cycle regulation, organelle organization, and neuronal differentiation (Fig. 3E), and showed binding similarity to neurodevelopmental transcription factors (Fig. 3F). In contrast, ZNF90-exclusive promoter-associated targets (465) did not show strong enrichment for specific functional categories.

We next focused on shared targets between OVOL2 and ZNF90. In total, 2,200 annotated genes were identified as shared targets, based on assignment to the nearest transcription start site (TSS). Shared peaks showed an even stronger depletion from intergenic regions compared to factor-specific peaks, accounting for only 18.84% and 25.93% of ZNF90 and OVOL2 peaks, respectively (Fig. 3D). This preferential localization to gene-associated regions supports a convergent role for both transcription factors in the direct regulation of shared transcriptional programs. Shared promoter-associated targets exhibited enrichment patterns like pioneer transcription factors such as FOXA1 and GATA3 (Fig. 3G), and included genes involved in cell cycle regulation (e.g., PRDM16, RUNX1). Notably, these were also the only annotated genes present in ZNF90-exclusive promoter peaks.

Although ZNF90-exclusive promoter-bound genes did not show strong global enrichment for specific functional categories, a subset of shared targets contained multiple peaks per gene (Fig. 3H). These included genes associated with extracellular matrix remodeling (*SMOC2*), neuronal maturation (*DPP6*, *DLGAP2*, *APBA2*, *ZNF536*), interneuron identity (*GABBR2*), progenitor cell cycle regulation (*ZNF423*, *HDAC4*, *SEPTIN9*), and ion channel function (*CACNA1C*, *CACNA1E*, *CACNA2D2*), many of which have been implicated in neurodevelopmental and neuropsychiatric disorders (Fig. 3H)^59–63^.

Finally, we asked whether the observed overlap in transcriptional targets reflects co-regulation within the same cells or regulation in distinct cellular populations. To address this, we examined the co-expression patterns of *ZNF90* and *OVOL2* in our previously published single-cell RNA-seq dataset of 30-day-old human COs^54^. Co-expression was highest in early proliferative aRG1 cells (∼2%; Fig. S6A) and, although overall low, was significantly higher than expected by chance across all aRG and bRG subtypes (Fig. S6B). These data suggest that *OVOL2* and *ZNF90* can act both cooperatively within the same cells and independently in distinct NPC subpopulations. Together, these findings indicate that OVOL2 and ZNF90 converge on gene networks controlling cell cycle progression and neurodevelopmental processes, while retaining distinct regulatory specificities.

### OVOL2 and ZNF90 induce distinct but partially overlapping transcriptional programs in rhesus macaque COs

To determine how OVOL2 and ZNF90 affect transcriptional programs, we ectopically expressed these transcription factors together with EGFP in 30-day-old rhesus macaque COs via electroporation^37^. Electroporated COs were cultured for an additional 4 days, thereby primarily targeting APs, with additional contribution from BPs^38^. COs were subsequently dissociated, EGFP-positive cells were isolated by FACS, and bulk RNA sequencing was performed (Fig. 4A, Fig. S7A; Materials and Methods).

**Figure 4.**
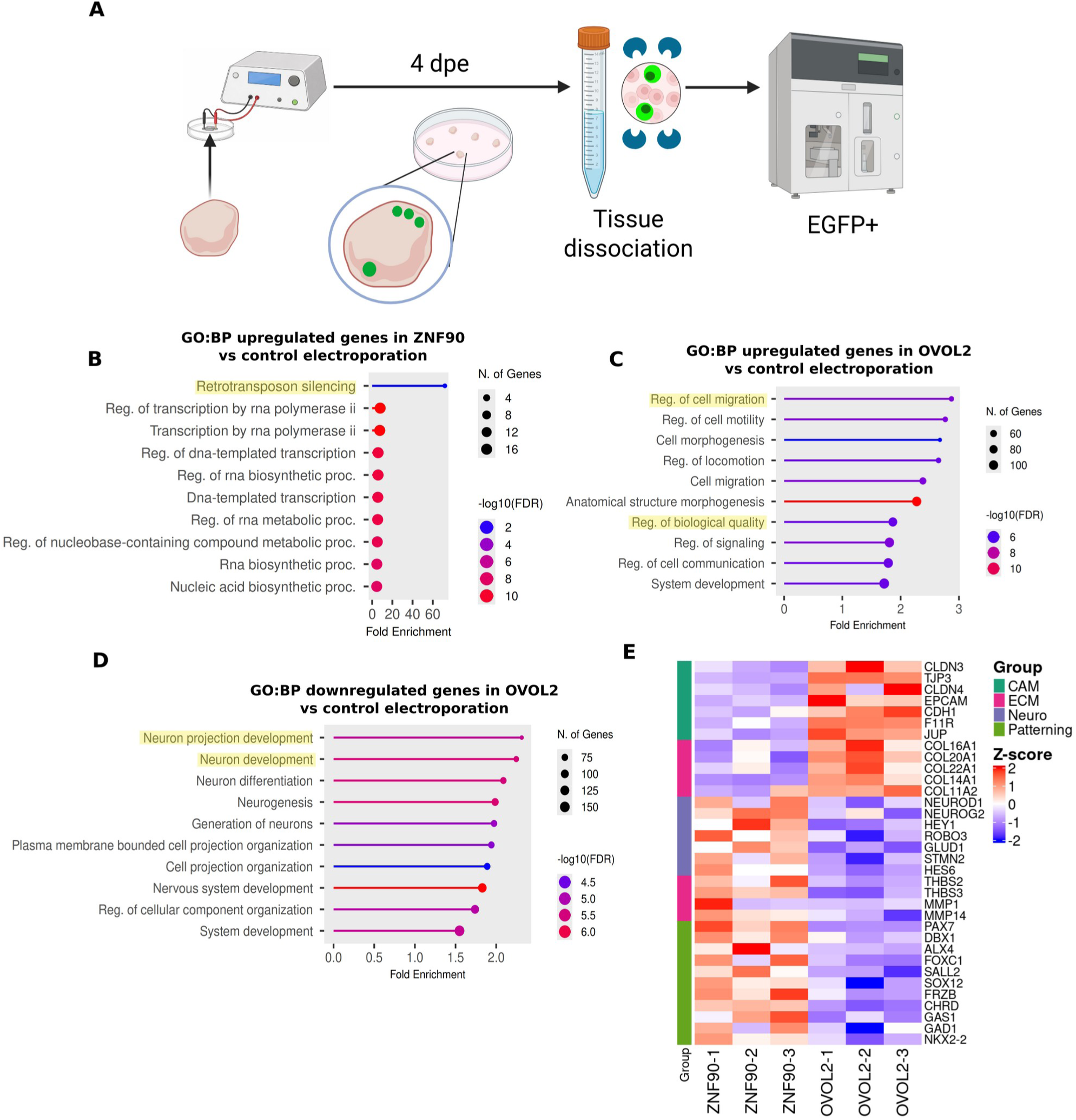
ZNF90 and OVOL2 elicit divergent but partially overlapping transcriptional responses following electroporation into rhesus macaque COs. (A) Schematic illustration of the experimental design to enrich electroporated GFP+ CO cells for transcriptome analysis. Key steps include electroporation of 30-day-old organoids, organoid dissociation and fluorescence-activated cell sorting (FACS) 4 dpe (n = 3 replicates per condition). (B) GO:BP enrichment analysis of DEGs (n = 38) between ZNF90-electroporated rhesus macaque CO cells and control electroporations. (C and D) GO:BP enrichment analysis of upregulated (C, n = 455) and downregulated (D, n = 595) genes in OVOL2-electroporated rhesus macaque CO cells. (E) Heatmap showing z-scored TPM values of selected DEGs involved in key biological processes (Group) between ZNF90 and OVOL2 electroporated CO cells.

Ectopic expression of ZNF90 resulted in a limited set of 38 differentially expressed genes (DEGs) (Table S7), which were nonetheless enriched for Gene Ontology biological processes related to retrotransposon silencing, chromosome segregation, and forebrain neuronal development (Fig. 4B, Table S8). In contrast, *OVOL2* ectopic expression induced a substantially broader transcriptional response, with 1,050 DEGs identified (Table S7), consistent with its markedly higher number of genome-wide binding sites compared to ZNF90. Functional enrichment analysis of the 455 upregulated genes revealed pathways associated with cell shape, migration, and regulation of biological quality (Fig. 4C, Fig. S7B), whereas the 595 downregulated genes were enriched for neuronal differentiation (Fig. 4D).

Comparative analysis of DEGs across both experimental conditions revealed distinct transcriptional programs. *OVOL2*-overexpressing cells showed enrichment for genes involved in cell adhesion molecules (CAMs) and extracellular matrix (ECM) deposition, whereas *ZNF90*-overexpressing cells exhibited increased expression of genes associated with neuronal differentiation, ECM remodeling, and dorsal telencephalic patterning (Fig. 4E). Despite these differences, both conditions shared a subset of DEGs linked to neurodevelopment, including *EMX1, DMRT3, GAL, FOXG1-AS1*, and cell cycle-associated genes such as *RP11-454P7.1* and *LINC01551* (Table S7).

Together, these results indicate that OVOL2 exerts a broad regulatory influence on transcriptional programs related to cell adhesion, morphology, and NPC state, whereas ZNF90 modulates a more restricted set of genes, including pathways linked to genome regulation and neuronal development, with partial convergence on shared neurodevelopmental targets.

### OVOL2 and ZNF90 promote AP maintenance at the expense of BPs and neurons

Next, we assessed the functional effects of *OVOL2* and *ZNF90* expression on NPCs behavior. To this end, we electroporated these genes individually and in combination into 30-day-old rhesus macaque COs, similar as previously described^39^. COs were analyzed 4 days post-electroporation (4 dpe; Fig. 5A), allowing the assessment of both APs and BPs. We first examined the distribution of electroporated (GFP-positive) cells within the VZ and SVZ/NL (Neuronal Layer). For OVOL2-, ZNF90-, and combined OVOL2 + ZNF90-electroporated COs, we observed a significant increase in the proportion of GFP-positive cells within the VZ compared to control (Fig. 5B).

**Figure 5.**
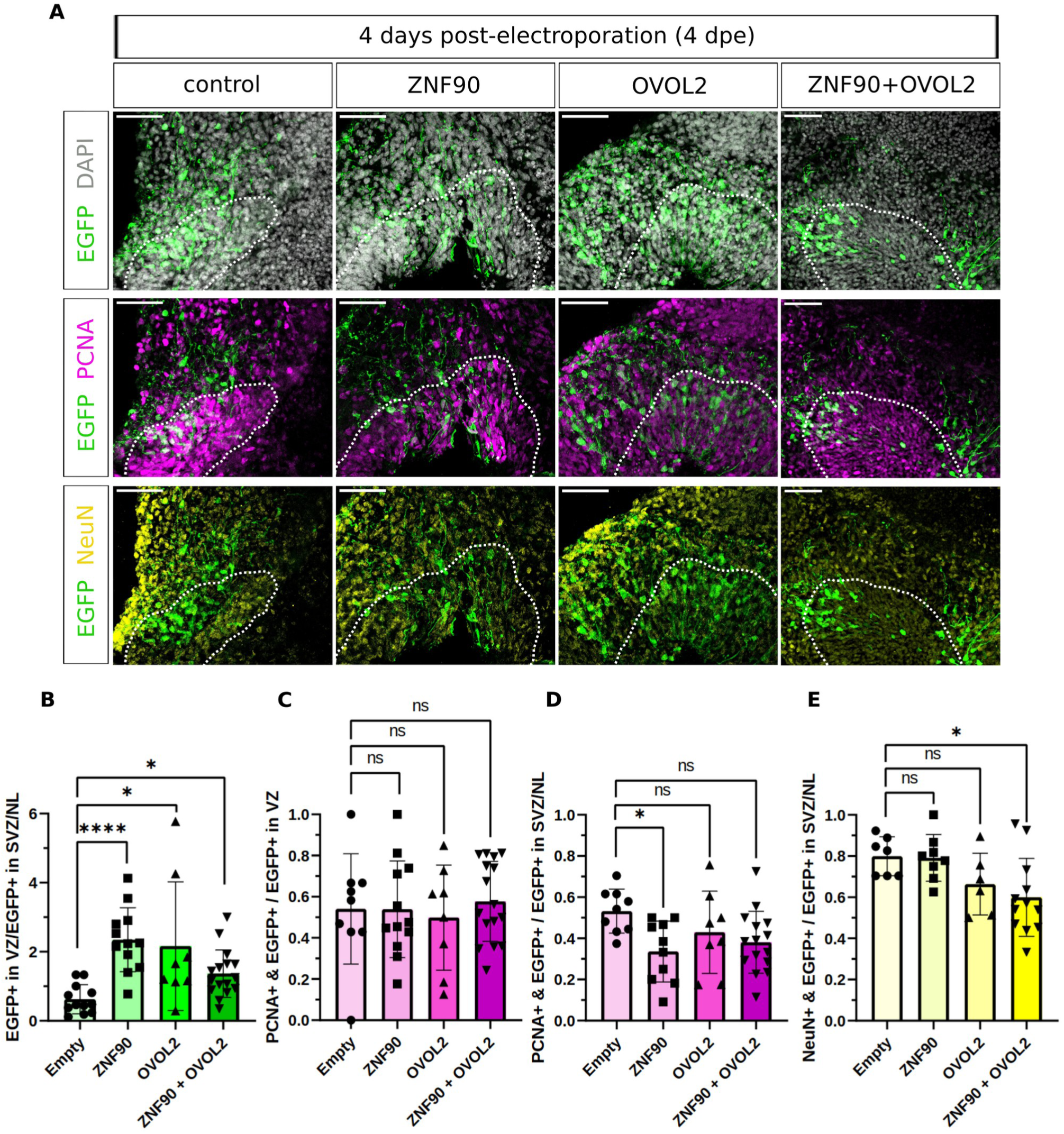
OVOL2 and ZNF90 transiently promote apical progenitor maintenance but differentially affect basal progenitors and neuronal output at 4 dpe. (A) Triple immunofluorescence for EGFP (green), PCNA (magenta), and NeuN (yellow), combined with DAPI (gray), in 34-day-old rhesus macaque COs at 4 dpe with EGFP plus either control, ZNF90, OVOL2, or ZNF90 plus OVOL2 (ZNF90+OVOL2) expression plasmids. Dashed lines indicate the basal boundary of the VZ. Scale bars, 50 μm. (B) Proportion of EGFP+ cells in the VZ to EGFP+ cells in the SVZ in 34-day-old rhesus macaque COs at 4 dpe. Data are the mean of 12 control, 12 ZNF90, 8 OVOL2, and 16 ZNF90+OVOL2 ventricle-like structures; error bars, ±SD; ns, not significant; *P < 0.05, ****P < 0.0001; Kruskal–Wallis test with Dunn’s correction. Exact adjusted P-values: Empty vs. OVOL2, P = 0.0148; Empty vs. ZNF90+OVOL2, P = 0.0379. (C and D) Proportion of PCNA+EGFP+ cells among EGFP+ cells within the VZ (C) and SVZ/NL (D) of 34-day-old rhesus macaque COs at 4 dpe. Data are the mean of control (9 and 9), ZNF90 (12 and 10), OVOL2 (8 and 8), and ZNF90+OVOL2 (16 and 15) ventricle-like structures for (C) and (D), respectively; error bars, ±SD; ns, not significant; *P < 0.05; one-way ANOVA with Dunnett’s correction. Exact adjusted P-values: (D) Empty vs. ZNF90, P = 0.0215. (E) Proportion of NeuN+EGFP+ cells among EGFP+ cells within the SVZ/NL of 34-day-old rhesus macaque COs at 4 dpe. Data are the mean of 7 control, 8 ZNF90, 6 OVOL2, and 12 ZNF90+OVOL2 ventricle-like structures; error bars, ±SD; ns, not significant; *P < 0.05; one-way ANOVA with Dunnett’s correction. Exact adjusted P-values: Empty vs. ZNF90+OVOL2, P = 0.0232.

We next asked whether this increase reflects elevated abundance of proliferative and/or mitotic cells in the VZ. However, we did not detect an increased proportion of PCNA-positive or mitotic (phospho-histone H3-, PH3-positive) cells within the VZ upon *OVOL2*, *ZNF90*, or combined expression (Fig. 5C, Fig. S8A,B). These data suggest that the accumulation of GFP-positive cells in the VZ is unlikely to result from increased abundance of proliferative and mitotic cells in the VZ but rather points to altered NPC dynamics favoring maintenance of the AP state which would affect basal cell populations.

To test this possibility, we analyzed basal cell populations, i.e. BPs and neurons. ZNF90 electroporation led to a significant reduction in the proportion of PCNA-positive cells in the SVZ/NL, indicative of reduced BP numbers. OVOL2 and combined OVOL2 + ZNF90 electroporation showed a similar trend, although not reaching statistical significance (Fig. 5D). Notably, this reduction was not accompanied by a decreased proportion of PH3-positive mitotic cells (Fig. S8A,B), suggesting that the lower BP abundance is not due to reduced mitotic activity of BPs. Analysis of neuronal (NeuN-positive) populations revealed no significant change upon *ZNF90* expression. In contrast, OVOL2 electroporation resulted in a reduction in neuronal proportions, which reached statistical significance in the combined OVOL2 + ZNF90 condition (Fig. 5E).

Together, these findings indicate that both OVOL2 and ZNF90 promote maintenance of APs, likely by limiting differentiation into BPs. OVOL2, alone and in combination with ZNF90, additionally reduces neuronal output, suggesting a broader effect on NPC maintenance and differentiation. These functional effects are consistent with the transcriptional programs and genomic binding profiles identified above, linking OVOL2 and ZNF90 to regulatory networks controlling NPC state and neurodevelopmental progression.

### OVOL2 and ZNF90 exert temporally distinct effects on NPC dynamics and neuronal output

To assess whether the observed effects on NPC dynamics are maintained over time, we analyzed electroporated rhesus macaque COs at 10 dpe (Fig. 6A). As at 4 dpe, we first examined the distribution of electroporated (GFP-positive) cells within the germinal zones (VZ and SVZ/NL). At 10 dpe, ZNF90- and combined OVOL2 + ZNF90-electroporated COs continued to show a significant increase in the proportion of GFP-positive cells within the VZ compared to control. In contrast, OVOL2-electroporated COs no longer displayed this enrichment (Fig. 6B), suggesting that the OVOL2-induced accumulation of APs is transient, whereas ZNF90-mediated effects are sustained.

**Figure 6.**
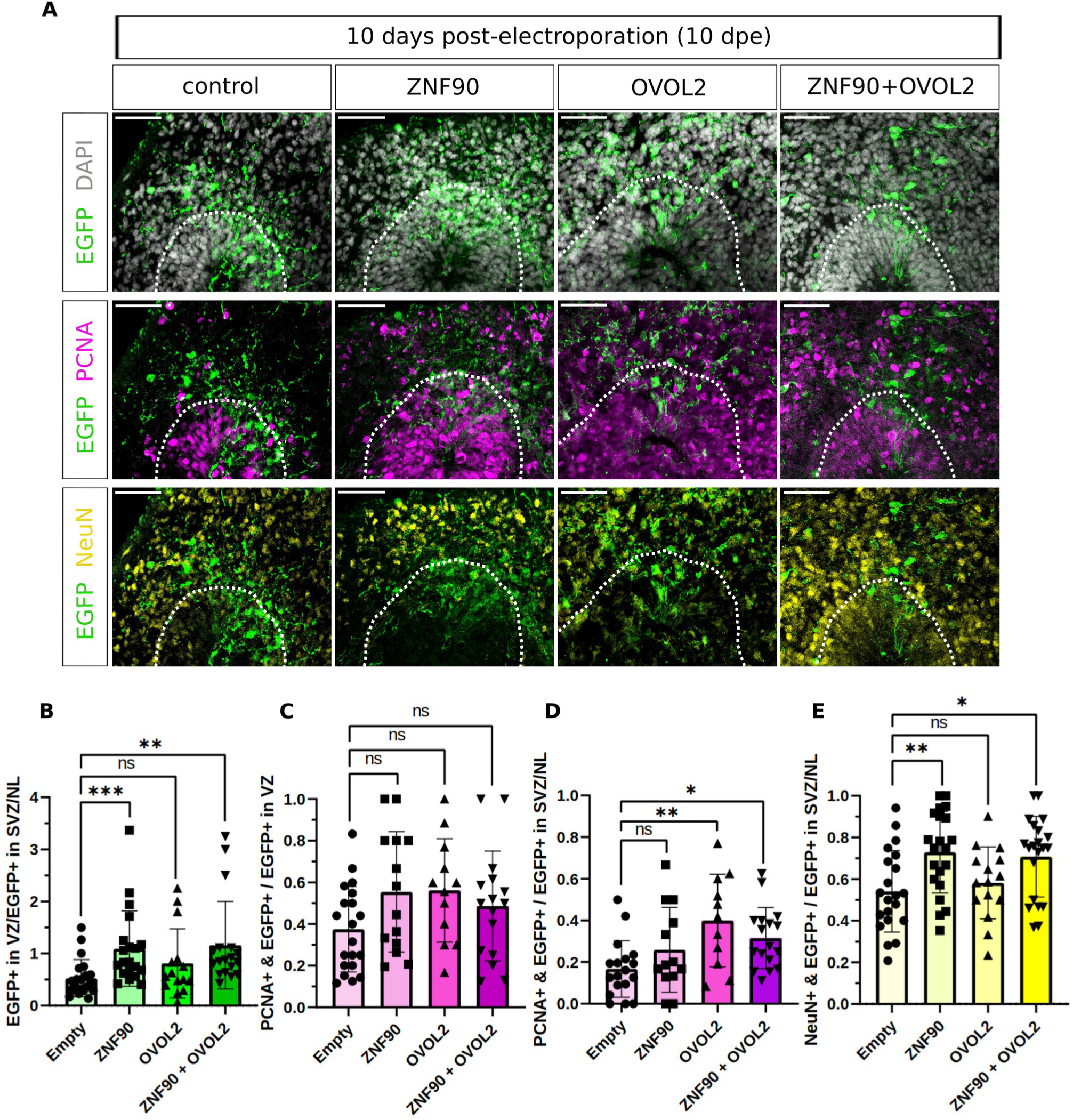
OVOL2 and ZNF90 exert temporally distinct effects on progenitor progression and neuronal output at 10 dpe. (A) Triple immunofluorescence for EGFP (green), PCNA (magenta), and NeuN (yellow), combined with DAPI (gray), in 40-day-old rhesus macaque COs at 10 dpe with EGFP plus either control, ZNF90, OVOL2, or ZNF90 plus OVOL2 (ZNF90+OVOL2) expression plasmids. Dashed lines indicate the basal boundary of the VZ. Scale bars, 50 μm. (B) Proportion of EGFP+ cells in the VZ to EGFP+ cells in the SVZ in 40-day-old rhesus macaque COs at 10 dpe. Data are the mean of 20 control, 19 ZNF90, 13 OVOL2, and 15 ZNF90+OVOL2 ventricle-like structures; error bars, ±SD; ns, not significant; **P < 0.01, ***P < 0.001; Kruskal–Wallis test with Dunn’s correction. Exact adjusted P-values: Empty vs. ZNF90, P = 0.0003; Empty vs. ZNF90+OVOL2, P = 0.0023. (C and D) Proportion of PCNA+EGFP+ cells among EGFP+ cells within the VZ (C) and SVZ/NL (D) of 40-day-old rhesus macaque COs at 10 dpe. Data are the mean of control (20 and 18), ZNF90 (14 and 13), OVOL2 (12 and 11), and ZNF90+OVOL2 (16 and 17) ventricle-like structures for (C) and (D), respectively; error bars, ±SD; ns, not significant; *P < 0.05, **P < 0.01; one-way ANOVA with Dunnett’s correction. Exact adjusted P-values: (D) Empty vs. OVOL2, P = 0.0027; Empty vs. ZNF90+OVOL2, P = 0.0396. (E) Proportion of NeuN+EGFP+ cells among EGFP+ cells within the SVZ/NL of 40-day-old rhesus macaque COs at 10 dpe. Data are the mean of 21 control, 21 ZNF90, 15 OVOL2, and 21 ZNF90+OVOL2 ventricle-like structures; error bars, ±SD; ns, not significant; *P < 0.05, **P < 0.01; one-way ANOVA with Dunnett’s correction. Exact adjusted P-values: Empty vs. ZNF90, P = 0.0058; Empty vs. ZNF90+OVOL2, P = 0.0166.

As at 4 dpe, we next assessed the proportion of proliferative and/or mitotic cells in the VZ. No increase in the proportion of PCNA-positive or PH3-positive cells was observed in the VZ under any condition (Fig. 6C, Fig. S8C,D), indicating that changes in VZ cell distribution in the ZNF90 condition are not driven by increased abundance of proliferative and mitotic cells in the VZ.

We then analyzed basal cell populations, i.e. BPs and neurons. At 10 dpe, ZNF90 electroporation did not result in significant changes in the proportion of PCNA-positive cells in the SVZ/NL compared to control. In contrast, OVOL2 and combined OVOL2 + ZNF90 electroporation led to a significant increase in PCNA-positive cells in the SVZ/NL (Fig. 6D), indicating an increased proportion of BPs at this stage. Notably, this increase was not accompanied by an increased proportion of PH3-positive mitotic cells (Fig. S8C,D), suggesting that the higher BP abundance is not due to elevated mitotic activity of BPs. Finally, we assessed neuronal (NeuN-positive) populations. OVOL2 electroporation did not significantly alter neuronal proportions. In contrast, ZNF90 and combined OVOL2 + ZNF90 electroporation resulted in a significant increase in NeuN-positive cells in the basal region (Fig. 6E). In addition, ZNF90 electroporation increased the proportion of NeuN-positive cells within the VZ (Fig. S8E), indicating that neuronal differentiation can occur while cells remain associated with the ventricular zone.

To determine whether these temporal differences are reflected in endogenous NPC populations, we analyzed previously generated single-cell RNA-seq data from 30-day-old human cerebral organoids^54^. ZNF90-positive cells displayed lower pseudotime values than OVOL2-positive or double-positive cells (Fig. S9A), consistent with a less advanced developmental state. Furthermore, proliferative activity in the self-renewing aRG1 population was not associated with expression of either gene. In contrast, the neurogenic aRG3 population exhibited increased proliferative signatures in ZNF90-positive (57.0%) and double-positive (64.4%) cells, whereas the self-renewing bRG1 population displayed higher proliferative activity in OVOL2-positive (84.5%) and double-positive (94.7%) cells (Fig. S9B).

Together, these findings reveal that OVOL2 and ZNF90 regulate NPC dynamics through distinct temporal mechanisms. OVOL2 promotes a transient stabilization of AP identity, delaying the transition to BPs and resulting in their subsequent accumulation at later developmental stages. In contrast, ZNF90 maintains AP representation over a prolonged period without suppressing neuronal differentiation, ultimately leading to increased neuronal output. Thus, whereas OVOL2 appears to primarily regulate the timing of apical-to-basal progenitor transitions, ZNF90 sustains AP characteristics while remaining permissive for neurogenesis. The increased abundance of NeuN-positive cells within the VZ following *ZNF90* expression is compatible with neuronal differentiation occurring within or in close association with the VZ. Collectively, these findings indicate that OVOL2 and ZNF90 do not simply promote AP maintenance but instead differentially regulate NPC dynamics through complementary mechanisms, thereby influencing the temporal balance between self-renewal, lineage progression and neuronal output.

## Discussion

In this study, we combined within-species and cross-species transcriptomic analyses to identify regulators of human NPC biology. By comparing human NPCs with both human neurons and corresponding rhesus macaque samples, we identified transcriptional programs enriched in human NPCs, including genes associated with NPC maintenance, VZ identity, proliferative capacity, and chromosome segregation. From these analyses, we prioritized two transcription factors with distinct evolutionary histories, the ape-specific zinc-finger protein ZNF90 and the deeply conserved transcription factor OVOL2^56^. Integrating transcriptomic profiling with CUT&RUN revealed that both factors regulate overlapping neurodevelopmental programs and share a subset of direct genomic targets, despite striking differences in the extent of their regulatory influence. Functional analyses in rhesus macaque COs demonstrated that both genes initially promote AP maintenance but subsequently drive distinct developmental trajectories: OVOL2 ultimately favors BP expansion, whereas ZNF90 promotes sustained AP maintenance accompanied by increased neuronal production. Co-expression of both factors combined key aspects of the individual phenotypes, leading to increased abundances of APs, BPs, and neurons. Together, these findings suggest that evolutionarily conserved and evolutionarily young transcription factors can converge on shared regulatory networks while exerting distinct effects on the balance between NPC maintenance and differentiation.

Several aspects of these findings warrant further discussion. First, our comparison of human and rhesus macaque NPCs revealed a coordinated enrichment of genes associated with VZ identity, NPC maintenance, and proliferative capacity in human NPCs. These included regulators of aRG organization, signaling pathways involved in NPC maintenance, mechanotransduction and cell adhesion, growth factor-mediated niche interactions, as well as genes required for DNA replication and cell-cycle progression. Collectively, these transcriptional differences are consistent with an enhanced capacity of human APs to maintain a proliferative state and expand the NPC pool. Such an expansion is thought to represent a key cellular foundation for the increased neuronal output that ultimately contributed to the evolutionary enlargement of the human neocortex^6,7,57^. Unexpectedly, however, we also identified a strong enrichment of genes involved in chromosome segregation and chromatid segregation fidelity among those enriched in human NPCs. This finding suggests that primate brain evolution was accompanied not only by mechanisms promoting NPC expansion but also by enhanced surveillance of genome integrity. Such a relationship is biologically plausible. As NPC proliferation increases, individual APs generate progressively larger clones through repeated rounds of cell division. Under these conditions, even small increases in chromosome segregation errors would be expected to have amplified consequences for tissue development. Improved regulation of genes associated with chromatid segregation may therefore represent an adaptation that supports larger NPC populations while preserving genomic stability. Notably, previous work suggested that increased chromosome segregation fidelity distinguishes modern humans from Neanderthals^64^. Our findings raise the possibility that evolutionary changes in chromosome segregation control extend beyond recent human evolution. Although the present study focuses on the functional characterization of OVOL2 and ZNF90, the enrichment of chromosome segregation programs in human NPCs identifies genome maintenance as an additional candidate mechanism that may have contributed to human neocortical evolution and warrants direct functional investigation.

Second, one of the principal findings of this study is that transcription factors with markedly different evolutionary histories become incorporated into overlapping developmental gene regulatory networks. Despite one factor being deeply conserved and the other being restricted to the ape lineage, OVOL2 and ZNF90 share genomic targets, regulate related transcriptional programs, and produce partially convergent cellular phenotypes. Both factors affected genes associated with neurodevelopment, NPC biology, and cell-cycle regulation, and they shared a subset of direct genomic targets. These findings suggest that OVOL2 and ZNF90 are incorporated into partially overlapping gene regulatory networks despite their markedly different evolutionary histories. This overlap was further reflected in the functional experiments, where ectopic expression of either factor initially promoted AP maintenance in rhesus macaque COs. However, their effects diverged at later developmental stages, resulting in distinct cellular outcomes. Whereas OVOL2 ultimately favored BP expansion, ZNF90 promoted sustained AP maintenance while simultaneously increasing neuronal production. Interestingly, the molecular footprints of the two transcription factors differed substantially. CUT&RUN identified 91,983 binding sites and transcriptomic analysis revealed 1,050 differentially expressed genes following OVOL2 expression, whereas ZNF90 was associated with only 6,599 binding sites and 38 differentially expressed genes. These observations are consistent with OVOL2 functioning as a broader transcriptional regulator, whereas ZNF90 may exert a more specialized and targeted regulatory role. One possible explanation is their contrasting evolutionary origins. *OVOL2* is deeply conserved across vertebrates^56^ and may therefore occupy a central position within established neurodevelopmental gene regulatory networks. In contrast, *ZNF90* emerged more recently during primate evolution^65,66^ and may not yet have achieved the same degree of network integration. Nevertheless, despite these differences, both factors converge on common neurodevelopmental targets and biological processes. This convergence suggests that evolutionary innovation can occur not only through the emergence of entirely novel pathways but also through the modification and refinement of existing regulatory networks by factors of distinct evolutionary ages.

Third, the functional effects of OVOL2 are consistent with its established role as a regulator of epithelial identity and antagonist of epithelial-to-mesenchymal transition (EMT)^48,49^. At early stages after electroporation, OVOL2 increased the proportion of APs, consistent with enhanced maintenance of VZ characteristics and reduced delamination. This interpretation is further supported by the identification of direct OVOL2 targets such as *ZEB1* and *ZEB2*, two central promoters of EMT that have previously been implicated in regulating NPC state transitions^57,67,68^. Whereas ZEB2 has been linked to promote transitions from neuroepithelial cells to radial glia^57^, our findings suggest that a similar regulatory logic may operate at later developmental stages by delaying the transition from APs to BPs through inhibition of *ZEB2* expression. Despite this early maintenance of APs, *OVOL2* expression subsequently resulted in an increased abundance of BPs. Several observations indicate that this increase likely originates from the initially expanded AP pool. While APs were elevated at 4 days post-electroporation, their abundance returned to control levels by 10 days post-electroporation, coinciding with the increase in BPs. At the same time, neuronal proportions remained largely unchanged, suggesting that many of these newly generated BPs had not yet differentiated further into neurons. The mechanisms underlying this delayed BP expansion remain unclear. It may reflect direct effects of the OVOL2-dependent gene regulatory network, compensatory responses within the enlarged AP population, intrinsic developmental trajectories of rhesus macaque COs, or a combination of these factors. From an evolutionary perspective, however, the net outcome would be a delayed apical-to-basal progenitor transition followed by expansion of the BP compartment, a process predicted to increase neuronal output.

In contrast, ZNF90 produced a distinct developmental trajectory. APs remained elevated at both 4 and 10 dpe, indicating sustained NPC maintenance. At 4 days post-electroporation, this was accompanied by a reduction in BPs, suggesting a transient inhibition of AP to BP transitions. By 10 dpe, BP abundance had returned to control levels despite the continued expansion of the AP pool, implying that the enlarged NPC population was able to restore BP production. The sustained maintenance of APs is also consistent with the proposed molecular functions of ZNF90. As a member of the KRAB zinc-finger protein family, ZNF90 has previously been linked to the repression of TEs, including human endogenous retroviruses (HERVs), which are increasingly recognized as important contributors to gene regulatory programs in NPCs^69^. Together with our transcriptomic and CUT&RUN analyses, these observations suggest that ZNF90 may contribute to NPC maintenance through the regulation of transposable element-associated regulatory networks that have become integrated into primate neurodevelopmental programs. Unexpectedly, *ZNF90* expression also increased neuronal abundance at later stages. This effect was accompanied by an increased proportion of NeuN-positive cells within the VZ, raising the possibility that ZNF90 promotes direct neurogenesis, the generation of neurons directly from APs without an intermediate BP stage. Several explanations may account for this observation. First, it could represent an organoid-specific phenomenon or a consequence of transgene overexpression. Second, it may reflect developmental stage-dependent shifts in neurogenic modes, as direct neurogenesis has been reported during later developmental windows^70^. Third, ZNF90 may influence the neurogenic behavior of BPs themselves, for example by promoting neurogenic divisions rather than NPC amplification, as previously reported for human^71^, and thereby increasing neuronal output without altering BP abundance. Although additional experiments will be required to distinguish between these possibilities, the transcriptional induction of extracellular matrix-remodeling metalloproteases and neurogenic regulators following *ZNF90* expression is consistent with a role in facilitating neuronal differentiation and lineage progression.

Finally, the partial overlap between OVOL2 and ZNF90 prompted us to investigate their combined activity. Both genes share direct targets, regulate overlapping sets of differentially expressed genes, and are co-expressed within a small subset of NPCs. Similar combinatorial approaches have previously revealed cooperative functions among evolutionarily young genes, including the metabolic interaction between the human-specific ARHGAP11B and the ape-specific factor GLUD2^72^, as well as the synergistic effects of the human-specific genes *NOTCH2NLB* and *NBPF14* on NPC expansion and delamination control^39^. Consistent with these precedents, co-electroporation of *OVOL2* and *ZNF90* initially promoted AP maintenance. At later stages, however, simultaneous expression increased the abundance of APs, BPs, and neurons, effectively combining key aspects of the individual phenotypes. Such a combined effect would be predicted to substantially enhance neurogenic output and could therefore have profound consequences for neocortical expansion. Although the endogenous co-expression of *OVOL2* and *ZNF90* is restricted to a relatively small NPC population, our findings provide evidence that evolutionary novel and evolutionarily conserved transcription factors can functionally interact within the same cellular context. Together, these findings suggest that evolutionary innovation does not depend solely on the emergence of new genes, but on their progressive incorporation into ancient developmental gene regulatory networks. Rather than acting independently, lineage-specific transcription factors can become integrated with conserved developmental regulators to generate novel combinations of NPC behaviors during primate corticogenesis.

## Resource Availability

### Lead contact

Michael Heide

### Materials availability

Requests for resources and reagents should be directed to M.H. and can be provided pending scientific review and a completed material transfer agreement.

### Data and code availability

Raw bulk RNA-seq and CUT&RUN-seq data generated in this study have been deposited in ENA under accession number PRJEB115502. Processed data, including gene expression matrices, R environments, and metadata used for downstream (re)analyses, are available in Zenodo (DOI: 10.5281/zenodo.21070930) for editorial review and will be made publicly accessible upon publication.

All custom scripts and computational workflows used for data preprocessing, (re)analysis, and figure generation are openly available at: https://github.com/MateoBastidasBetancourt/Bastidas_Betancourt_et_al_2026

## Acknowledgements

We apologize to all researchers whose work could not be cited due to space limitations. We thank G. Salinas and the team at the NGS Integrative Genomics (NIG) facility at the University Medical Center Göttingen (UMG), as well as the Gesellschaft für wissenschaftliche Datenverarbeitung mbH Göttingen (GWDG), for RNA-seq and CUT&RUN sequencing services and bioinformatic infrastructure. We are grateful to R. Behr for access to personnel and infrastructure and for providing the rhesus macaque iPSC line, to Dr. I. Niehaus for providing the human iPSC line, and to Dr. Ranjit Pradhan for providing the cDNA samples of human neuronal culture. We also thank the members of the Heide lab for helpful discussions and critical reading of the manuscript. MH was supported by an ERC starting grant (PRIMAZINC, 101039421).

## Author contributions

Conceptualization: CMB, MH Methodology: CMB Software: CMB

Investigation: CMB, IN, IBG, DB, MPS, AMC, MR, SH

Visualization: CMB, IN, IBG, DB, MPS, AMC, MR

Resources: MI, LV, TK, EK, AT, SH

Formal analysis: CMB, IN, IBG, DB, MPS, AMC, MR

Data curation: CMB

Supervision: MH, AT

Project administration: MH

Funding acquisition: MH

Writing—original draft: CMB, MH

Writing—review & editing: CMB, MH

## Declaration of generative AI and AI-assisted technologies

During the preparation of this work, the authors used the OpenAI GPT-5.5 service provided by GWDG to assist with language editing, grammar and spelling correction, and code troubleshooting and formatting. All AI-assisted output was reviewed and edited by the authors, who take full responsibility for the final content of the manuscript.

## Supplemental information

### Document S1. Figures S1–S9 and Tables S9-S10

Table S1. Metadata of the previously published RNA-seq samples of human and rhesus macaque cortical development that were analyzed in this study.

Table S2. Differentially expressed genes identified between human NPCs and neurons.

Table S3. Genes with human- or rhesus-biased NPC enrichment in the pooled datasets.

Table S4. HOMER known-motif analysis output of ZNF90-bound regions.

Table S5. ZNF90 and OVOL2 annotated peaks to genomic features and closest gene TSS.

Table S6. ZNF90 and OVOL2 peaks overlapping annotated repeats in the corresponding genome.

Table S7. Differentially expressed genes between the control, ZNF90 and OVOL2 electroporations.

Table S8. Gene set enrichment analysis output of the Log2FC-sorted genes resulting from the ZNF90 vs control comparison.

## Methods

### Ethics

Human PCW 14 fetal brain tissue was obtained from spontaneous abortions at the Hospital for Obstetrics and Gynecology “Prof. Dimitar Stamatov,” Medical University of Varna, Bulgaria, following written informed maternal consent and approval by the local Ethics Committee (Protocol No. 19/April 2012 and Protocol No. 55/June 2016). The *in situ* hybridization experiment was conducted in accordance with Resolution No. 19 of the Commission on Ethics in Scientific Research at the Medical University of Varna (5 April 2012), including subsequent amendments (Resolution No. 66, 26 August 2019; Resolution No. 141, 14 March 2024). Gestational age was determined either by Crown-Rump Length (CRL) or based on the date of the last menstrual period reported by the mother.

Following collection, fetal heads were fixed in 4% paraformaldehyde (PFA) in phosphate-buffered saline (PBS; pH 7.5) for 24 h. Brains were then dissected, post-fixed in the same fixative for 5–7 days, cryoprotected in sucrose, embedded in optimal cutting temperature (OCT) compound, and cryosectioned at 20 μm thickness.

### Cell lines

#### Induced Pluripotent Stem Cells (iPSCs)

One male rhesus macaque iPSC (iRh33.1) line was used in this study and cultured according to the original protocol (Stauske et al., 2020)^73^. Cells were maintained at 37 °C and 5% CO_2_ in iPSB+ medium (StemMACS™ iPS-Brew XF supplemented with StemMACS™ iPS-Brew XF 50x supplement, 1 μM IWR-1, and 0.5 μM Chir99021) on Geltrex-coated dishes (0.167 mg/mL). The human iPSC line CRTDi011-A^74^ was cultured in mTeSR1 (Stemcell Technologies) on Geltrex-coated dishes as previously described. For both cell lines, the culture medium was changed daily, and cells were passaged at 70-80% confluency using Versene for iRh33.1 and ReLeSR (Stemcell Technologies) for CRTDi011-A.

#### Neural Progenitor Cells (NPCs)

Human NPCs were represented by the immortalized ReNcell® CX NPC line (Merck)^52^. Cells were maintained in ReNcell® NSC Maintenance Media (Merck Millipore) supplemented with FGF-2 (20 ng/mL) and EGF (20 ng/mL) on laminin-coated dishes (20 μg/mL in DMEM/F12 without HEPES, with L-glutamine; Merck Millipore). Cells were passaged using Accutase cell detachment solution (Merck Millipore) according to the manufacturer’s instructions. Unless otherwise stated, RNA was extracted at approximately 80% confluency.

### Generation of Rhesus Macaque and Human COs

COs were differentiated as previously described^37,74^. Briefly, 9000 cells per well were seeded in an ultra-low attachment 96-well plate (Thermo Fisher Scientific) in the respective iPSC medium supplemented with 50 μM ROCKi (Merck) to generate embryoid bodies (EBs). After 48 hours of incubation, the culture medium was changed to the respective medium without ROCKi. On day 5 post-seeding, the medium was changed to neural induction medium [DMEM/F12 (Thermo Fisher Scientific) containing 1% N2 supplement (Thermo Fisher Scientific), 1% GlutaMAX supplement (Thermo Fisher Scientific), 1% MEM nonessential amino acids (Thermo Fisher Scientific), and heparin (1 μg/ml; Sigma-Aldrich)] and changed every 48 hours. On day 7 (rhesus macaque) and day 9 (human) post-seeding, the EBs were embedded in Matrigel (Corning) and further cultured in differentiation medium [1:1 DMEM/F12 (Thermo Fisher Scientific)/neurobasal (Thermo Fisher Scientific) containing 1% B27 supplement without vitamin A (Thermo Fisher Scientific), 0.5% N2 supplement (Thermo Fisher Scientif- ic), 1% GlutaMAX supplement (Thermo Fisher Scientific), 0.5% MEM nonessential amino acids (Thermo Fisher Scientific), 1% penicillin-streptomycin (Thermo Fisher Scientific), 0.025% insulin solution (Sigma-Aldrich), and 0.00035% 2-mercaptoethanol (Merck)] on an orbital shaker with medium changes every 48 hours. 6 days after embedding, the medium was switched to differentiation medium containing B27 supplement with vitamin A (Thermo Fisher Scientific) and changed every 3 to 4 days. COs were grown on an orbital shaker in a 37°C incubator with a humidified atmosphere of 5% CO_2_ and 95% air.

### Rhesus Macaque and Human *in vitro* differentiated NPC and neuron culture

Rhesus macaque NPCs were differentiated from iRh33.1 iPSCs using the STEMdiff™ SMADi Neural Induction Kit (Stemcell Technologies) following the EB culture protocol. Briefly, EBs were generated as described before, with daily medium changes performed using STEMdiff™ Neural Induction Medium + SMADi. On day 5, EBs were replated on Matrigel-coated dishes (0,0013%) and further cultured with daily medium changes. On day 12, neural rosettes were replated using STEMdiff™ Neural Rosette Selection Reagent (Stemcell Technologies), Accutase-passaged twice, and cultured in STEMdiff™ Neural Progenitor Medium (Stemcell Technologies). For neuronal differentiation, RNA samples were kindly provided by Dr. Ranjit Pradhan (DZNE, Göttingen) and used in this study for RT-qPCR analysis. Samples were generated according to previously published protocols^75,76^.

### RNA extraction, cDNA synthesis, RT-qPCR

COs and cell pellets were dissociated in lysis buffer and processed for RNA isolation using the RNeasy® Micro Kit (QIAGEN), according to the manufacturer’s protocol, including DNase digestion. RNA was eluted in RNase-free water, and 0.1–1 µg were used for the cDNA synthesis reaction, according to the Maxima First Strand cDNA Synthesis Kit for RT-qPCR (Thermo Fisher Scientific).

To assess the temporal expression levels of *ZNF90*, *OVOL2*, and *ZEB2* (well-known target of OVOL2) in human NPCs, RT-qPCR was performed throughout development of human and rhesus (only for *OVOL2* and *ZEB2*) COs, as well as *in vitro* differentiated NPCs (see above) and neurons. Primers (Table S9) were designed in Primer3 using standard parameters^77^. Three replicates per time point were used for RT-qPCR. Reactions were prepared in MicroAmp™ Fast Optical 96-Well Reaction Plates (Applied Biosystems) and run on a StepOnePlus™ Real-Time PCR System. To calculate the relative expression of each gene, the delta-delta CT method was used^78^ and normalized to β-Actin as housekeeping gene.

### Generation of overexpression vectors

The coding sequences (CDS) of human *ZNF90* and rhesus macaque *OVOL2* were amplified from cDNA using PrimeStar Max DNA Polymerase (Takara Bio) under cycling conditions optimized according to amplicon length. Forward primers included a Kozak sequence and were designed according to the manufacturer’s requirements for In-Fusion cloning (Takara Bio), with homology arms to XhoI restriction enzyme site of linearized pCAGGS vector^79^. The NucleoSpin Gel and PCR Clean-up Kit (Macherey-Nagel) was used for gel purification. Transformation was performed in Stellar competent cells (Takara Bio), and plasmid DNA was isolated using the GeneJET Plasmid Miniprep Kit (Thermo Fisher Scientific) and the PureLink HiPure Plasmid Filter Maxiprep Kit (Thermo Fisher Scientific), following the manufacturers’ protocols. Furthermore, 5′-2xHA epitope-tagged versions of both CDS were cloned for CUT&RUN as described above, using two PCR reactions based on the previously cloned pCAGGS expression vectors. Primers are listed in Table S9.

### Microinjection and electroporation of rhesus macaque and human COs

Electroporation of COs was performed as previously described^37,38^. The following constructs were microinjected: pCAGGS-based constructs encoding *ZNF90* or *OVOL2*, either untagged or fused to an N-terminal 5′-2×HA epitope tag (pCAGGS-ZNF90, pCAGGS-OVOL2, and pCAGGS-OVOL2-5′-2×HA) in 30-day-old rhesus macaque COs, and pCAGGS-ZNF90-5′-2×HA in 30-day-old human COs. All constructs were co-electroporated with pCAGGS-EGFP as previously demonstrated^38^, except for CUT&RUN experiments.

Briefly, three to five ventricle-like structures per 30-day-old CO were microinjected with the electroporation mix containing the expression plasmids, according to Table S10, combined with 0.1% Fast Green dye (Carl Roth, 0301.2) in PBS. Microinjected COs were transferred to an in-house-built Petri dish electrode chamber connected to a square-wave electroporator (ECM 830, BTX) and covered with a small amount of DMEM/F12. Electroporation was performed by applying five pulses of 80 V, with a pulse duration of 50 ms and an interval of 1 s, to the microinjected COs. COs were then transferred to sterile 35-mm cell culture dishes filled with pre-warmed differentiation medium containing vitamin A and placed on an orbital shaker at 55 rpm in an incubator at 37 °C in a humidified atmosphere of 5% CO₂ and 95% air. Electroporated COs were further cultured for 4 or 10 days before fixation, or for 4 days before tissue dissociation, with medium changes every 3 days.

### Dissociation of COs for FACS and CUT&RUN

For FACS enrichment of GFP+ cells and CUT&RUN, 150–300 mg of tissue sample, corresponding to 5–10 electroporated COs, were dissociated using a trypsin-based Neural Tissue Dissociation Kit (Miltenyi Biotec) according to the manufacturer’s protocol. COs were bisected and gently agitated in ice-cold Hanks’ Balanced Salt Solution (HBSS) to remove dead cells from the inner core. The COs were then incubated with the enzymes indicated in the manufacturer’s protocol, stained with Trypan Blue, and counted using a hemocytometer. This yielded 1–5 million cells/sample collected in FACS buffer (PBS supplemented with 2% fetal bovine serum and 1 mM EDTA) for FACS, and 500,000 cells/sample collected in wash buffer (20 mM HEPES, 150 mM NaCl, 0.5 mM spermidine, 1x protease inhibitor) for CUT&RUN.

### FACS enrichment of EGFP+ cells

Cell suspensions were passed through a 70 μm strainer immediately before sorting. FACS was performed on a Sony SH800 sorter with the fluorescent protein filter set, using a 100 μm chip (LE-C3213, Sony). Cells were sorted into 500 µL of lysis buffer (Micro RNeasy kit, Qiagen) prior to RNA extraction, as described above.

### Bulk RNA-seq pre-processing and mapping

For bulk RNA-seq of electroporated rhesus macaque COs, raw sequencing signals were converted to BCL files using Illumina BaseCaller, followed by demultiplexing into FASTQ files with bcl2fastq v2.20.0.422. Samples from published human^16,18,40,41^ and rhesus macaque^43–45^ bulk transcriptome datasets of *in vivo* and *in vitro* NPC and neuronal samples are described in Table S1 and were analyzed together as described below.

Read quality assessment and adapter trimming were performed for all samples using Cutadapt^80^ and FastQC^81^. Reads were then mapped using STAR^82^ with the default alignment settings. Human reads were aligned to the GRCh38 genome, and rhesus macaque reads were aligned to the previously published human-to-rhesus lifted-off RheMac10 genome annotation^46^. Gene-level counts were generated using STAR’s --quantMode GeneCounts option.

### CUT&RUN and library preparation

CUT&RUN was performed on dissociated COs electroporated with either ZNF90-HA (N-terminal 5′-2×HA-tagged pCAGGS-ZNF90), ZNF90 (pCAGGS-ZNF90), OVOL2-HA (N-terminal 5′-2×HA-tagged pCAGGS-OVOL2), or OVOL2 (pCAGGS-OVOL2) (see “Electroporation of rhesus macaque and human COs”) using the CUTANA CUT&RUN Kit v5 (Epicypher, 14-1048), with the K-MetStat Panel (Epicypher, 19-1002) and E. coli spike-in DNA (Epicypher, 18-1401), according to the manufacturer’s protocol. Briefly, concanavalin A-conjugated paramagnetic beads were activated, and cells were resuspended in wash buffer before being adsorbed to the beads and permeabilized with digitonin (0.01%). Bead-bound cells were incubated overnight at 4 °C under constant nutation with either anti-HA tag antibody (Epicypher, 13-2010, 1:50, RRID: AB_3094663) or anti-rabbit IgG antibody (Epicypher, 13-0042, 1:50, RRID: AB_2923178). One anti-HA and one anti-IgG sample were included for each treatment. The next day, samples were washed and incubated with pAG-MNase and MgCl_2_, with wash steps in between, before stopping cleavage, resuspending the samples, and isolating antibody-bound DNA fragments.

DNA amounts were assessed using Qubit, and library preparation was performed with the CUTANA™ CUT&RUN Library Prep Kit (Epicypher, 14-1001), starting from 10 ng of DNA. The size distribution of the final DNA libraries was determined with the SS NGS Fragment 1–6000 bp Kit on the Fragment Analyzer, yielding an average fragment size of 340 bp. Libraries were sequenced on a NovaSeq 6000 using an S2 flow cell (100 cycles), generating approximately 10 million reads per CUT&RUN sample.

### ChIP/CUT&RUN-seq data processing and peak annotation

CUT&RUN-seq reads were pre-processed as described above for bulk-RNAseq. Reads were aligned to combined host–E. coli reference genomes (human hg38 + E. coli or rhesus macaque rheMac10 + E. coli) using Bowtie2 (v2.5.4)^83^ in very-sensitive mode with dovetail alignment enabled. Reference indices were pre-built using Bowtie2-build. Alignments were converted to BAM format, sorted, and indexed using Samtools (v1.21)^84^.

For downstream analysis, PCR duplicates were removed. Reads mapping to the E. coli genome were separated from host-mapped reads to generate organism-specific BAM files, which were subsequently indexed for deepTools bamCoverage (v3.5.6)^85^ track generation and MACS3 (v3.0.3)^86^ peak calling using standard parameters. For each TF, two peak-calling strategies were applied: (i) using the corresponding HA-IgG sample and (ii) using Samtools-generated merged untagged samples as controls (v1.21)^84^. High-confidence peaks were defined as the intersection of peaks identified in both analyses using bedtools (v2.31.1)^87^. Motif enrichment analysis and peak annotation to the hg38 and rheMac10 reference genomes were performed using HOMER (v5.1) ^88^. To assess overlap with repetitive elements, peak coordinates were intersected with RepeatMasker annotations downloaded from UCSC^,89,90^ using bedtools (v2.31.1)^87^. Overlap between target gene sets was calculated after assigning each peak to its closest TSS.

For the reanalysis of previously published ChIP-seq data from human and rhesus macaque PCW 5–12 telencephalic H3K4me2 and H3K27ac datasets^53^, processed files containing region coordinates were assigned to annotated gene promoters (+/− 3 kbp from the TSS) and intronic regions of the hg19 and rheMac2 genomes, respectively. Regions were considered reproducible if they showed a minimum overlap of 1 bp between two ChIP-seq replicates from the same time point and species, according to the original analysis^53^. In addition, replicated H3K27ac ChIP-seq peaks from day-30-old human COs (n = 3 isogenic replicates with 2 technical replicates each) and *in vitro* differentiated NPCs (n=3 isogenic replicates)^41^ were uploaded to the UCSC Genome Browser hub^89^ under the following sharing link: https://www.genome.ucsc.edu/s/cbastid/H3K27ac%20in%20human%20NPCs%20and%20d30%20organoids

### Phylogenetic analysis of *ZNF90* and the *OVOL2* core promoter

To investigate the evolutionary emergence of *ZNF90*, the ten closest paralogous protein sequences, as assessed by BLAST score, were identified from Ensembl annotations based on the human reference genome GRCh38. Protein sequences were retrieved and aligned using MUSCLE (v3.8)^91^ with default parameters. The core promoter sequences of *OVOL2* orthologs from human, chimpanzee, gorilla, and rhesus macaque were aligned using the same approach.

Phylogenetic analyses were performed within the UGENE platform (v48.0)^92^, implementing the PhyML maximum likelihood framework (v3.0)^93^. Trees were reconstructed under the LG amino acid substitution model, with simultaneous optimization of tree topology, branch lengths, and substitution rates. Branch support was estimated using the SH-like approximate likelihood ratio test^94^. Tree space was further explored using a combination of subtree pruning and regrafting (SPR) and nearest neighbor interchange (NNI) to refine topology and maximize likelihood.

To complement sequence-based phylogenetic inference, structural conservation among *ZNF90* paralogs was assessed using predicted three-dimensional models retrieved from the AlphaFold Protein Structure Database^95,96^ via the Protein Data Bank interface. Structural multiple alignments were performed using the US-align algorithm^97^, enabling quantitative comparison of global fold similarity across paralogs.

### Riboprobe synthesis and in-situ hybridization (ISH)

Total RNA was isolated from 50 mg of embryonic cortical tissue pooled from three specimens using the RNeasy Mini Kit (QIAGEN) and reverse-transcribed into cDNA using SuperScript III Reverse Transcriptase (Thermo Fisher Scientific) with oligo(dT) primers. Gene-specific primers containing T7 and SP6 promoter sequences were designed using Primer3 and used for PCR amplification of target fragments, which were subsequently sequence-verified. DIG-labelled antisense riboprobes were synthesized using SP6 RNA polymerase (New England Biolabs) and DIG RNA Labelling Mix (Roche Diagnostics). Following DNase treatment and ethanol/ammonium acetate precipitation, probes were resuspended in hybridization buffer (Ambion) at a concentration of 100 ng/µL and stored at −20 °C.

For colorimetric ISH, sections were rehydrated, treated with HCl and proteinase K, and hybridized overnight at 60 °C with 500–1500 ng of DIG-labelled probe. Following post-hybridization stringency washes, probe binding was detected using an anti-DIG alkaline phosphatase-conjugated antibody (Roche Diagnostics, Germany, RRID: AB_514497) and visualized with NBT/BCIP substrate (Sigma Aldrich). Sections were mounted with Hydro-Mount (Micro Tech Lab, Austria) and scanned with an AxioImager Z.2 microscope.

### Immunofluorescence

COs were fixed in 4% PFA in DPBS (pH 7.5) for 15 min to 1 h at room temperature. Following fixation, COs were washed three times in DPBS (Gibco, Thermo Fisher Scientific™) and stored in DPBS at 4°C until further processing. In preparation for cryosectioning, COs were first sequentially incubated in 15% and 30% sucrose in DPBS overnight at 4°C, then embedded in cryomolds filled with TissueTek OCT (Sakura) and frozen on dry ice. Cryosections of 20 µm thickness were obtained using a CryoStar NX70 cryostat (Thermo Fisher Scientific), mounted onto SuperFrost Plus slides (Epredia), and stored at −20°C. Immunofluorescence staining was performed as previously described^38,39^ In brief, sections were thawed and rinsed in PBS (pH 7.4). Antigen retrieval was performed in 0.1 M sodium citrate buffer (pH 6.0) for 1 h at 70°C, followed by cooling to room temperature. After washing in PBS, sections were permeabilized in 0.3% Triton-X100 in PBS (pH 7.4) and subsequently incubated in 0.1 M glycine in PBS (pH 7.4) for 30 min each. Blocking was carried out using 15% FCS in PBS for 30 min. Primary antibodies were diluted 1:200–1:300 in Can Get Signal® immunostain solution B (Toyobo) and applied overnight at 4°C. The following primary antibodies were used: PCNA (mouse monoclonal, CBL407, Merck, RRID: AB_93501), NeuN (rabbit polyclonal, ab104225, Abcam, RRID: AB_10711153), GFP (chicken, GFP-1020, Aves labs, RRID: AB_10000240) and PH3 (rat monoclonal, ab10543, Abcam, RRID: AB_2295065). The slides were washed in 15% FCS in PBS and then incubated with secondary antibodies diluted 1:500 in Can Get Signal solution B (Toyobo) together with DAPI (1:1000) for 1 h at room temperature. The following secondary antibodies were used: Alexa Fluor 488 anti-chicken (goat polyclonal, A-11039, Thermo Fisher Scientific, RRID: AB_142924), Alexa Fluor 555 anti-rabbit (donkey polyclonal, A-31572, Thermo Fisher Scientific, RRID: AB_162543), and Alexa Fluor 647 anti-mouse (donkey polyclonal, A-31571, Thermo Fisher Scientific, RRID: AB_162542). Slides were mounted with Mowiol (Carl Roth) and stored at 4°C in the dark.

### Image acquisition

Confocal images of immunostained sections were collected using a Zeiss LSM 800 microscope operated through ZEN Blue software and equipped with 10×/0.45 M27 and 20×/0.75 air objectives. For larger fields of view captured as tile scans, individual tiles were automatically stitched using Zeiss ZEN software. For Z stack acquisition, images were typically acquired at step intervals of 3.24 µm with the 10× objective and 2 µm with the 20× objective.

### Immunofluorescence- and image-based analyses

#### VZ to SVZ/NL Boundary

The boundary between the VZ and the SVZ/NL was identified based on differences in radial organization, nuclear density and PCNA immunofluorescence, which shows a higher intensity that is typical for the VZ.

#### Cell Type Quantifications

All cell quantifications of rhesus macaque COs were performed blinded using at least two independent CO batches. Cell counts were performed in ZEN Lite. First, EGFP-positive cells were counted in the VZ and SVZ/NL areas of each electroporated ventricle-like structure. The number of double-positive cells for PCNA, NeuN, or PH3 together with EGFP was then counted separately for each compartment. Data obtained from electroporated samples are represented as proportions of the EGFP-positive cell population. To avoid pseudo-replication, approximately 10 COs per condition and time point were imaged, with approximately one good-quality electroporated ventricle-like structure imaged per CO.

#### Statistical Analyses

All statistical analyses were performed in Prism (GraphPad Software). Outliers were identified using the ROUT method (Q = 1%). Normality of the data was evaluated using the Anderson–Darling, D’Agostino–Pearson, Shapiro–Wilk, and Kolmogorov–Smirnov tests. For comparisons between two groups, either an unpaired t-test or a Mann–Whitney test was applied depending on data distribution. For analyses involving more than two non-normally distributed groups, a Kruskal–Wallis test followed by Dunn’s multiple-comparison test was used. For analyses involving more than two normally distributed groups, one-way ANOVA followed by Dunnett’s multiple-comparison test was used. Data are presented as mean ± SD. Statistical significance was defined as P < 0.05. Exact adjusted P-values and sample sizes are reported in the corresponding figure legends and refer to the number of analyzed ventricle-like structures.

### Bioinformatic analyses

#### Bulk RNA-seq Analysis

Differential expression analysis was conducted using the edgeR package (v3.42.4)^98^ with library size normalization and removal of lowly expressed genes. For the analysis of previously published datasets, genes were considered differentially expressed (human NPC vs human neurons or human NPC–human neuron vs rhesus NPC–rhesus neuron) if they showed an absolute log2 fold change (Log2FC) > 1 and a false discovery rate (FDR) < 0.01. Gene ontology (GO) functional enrichment analysis of differentially expressed genes was performed using standard ShinyGO (v0.76.3)^99^ and GSEA^100^ parameters. Volcano plots of candidate genes were generated using VolcaNoseR^101^. HAR genomic coordinates were obtained from Whalen et al., 2023^42^. For each upregulated, downregulated, or non-differentially expressed (non-DE) gene from the human NPC vs human neuron differential expression analysis, the genomic distance to the nearest HAR was calculated. Genes were classified as HAR-proximal if located within 100 kb of the nearest HAR, based on the well-established cis-regulatory functions of HARs^102^. The proportion of HAR-proximal genes was computed for each group, and pairwise statistical differences between groups were assessed using Fisher’s exact test.

To assess the biological relevance of human NPC-upregulated genes, a WGCNA network was constructed from the combined human and rhesus macaque datasets using the WGCNA package^103^ with a soft-thresholding power of 5. Gene modules were identified by hierarchical clustering (minModuleSize = 30, deepSplit = 2) and merged at a cut height of 0.25. Modules associated with human NPC samples, based on module eigengene expression patterns, were selected for downstream analysis. Hub genes were defined based on intramodular connectivity, and functional enrichment of module genes was assessed using GO analysis as previously described. Bulk RNA-seq of electroporated rhesus macaque COs was analyzed using the same workflow, except that the Log2FC and FDR thresholds were set to 0.5 and 0.05, respectively, due to sample size differences relative to the cross-species analysis.

For the RT-qPCR data, three independent biological replicates were collected for each sample. Statistical significance for each comparison was assessed using a two-tailed unpaired Student’s t-test.

#### CUT&RUN-seq Enrichment

Binding enrichment across genomic features and TEs was assessed using Fisher’s exact test on 2 × 2 contingency tables, followed by Benjamini–Hochberg FDR correction for multiple testing. Motif enrichment significance was calculated by HOMER (v5.1)^88^ using its standard motif enrichment framework with Benjamini–Hochberg multiple testing correction. TF enrichment results for promoters bound by ZNF90, OVOL2, or both were generated using Enrichr’s internal databases^104^ with Benjamini–Hochberg multiple testing correction.

#### scRNA-seq Re-analysis Of Human And Marmoset COs

To dissect the expression of both genes in our COs, the Tynianskaia et al., 2026 scRNA-seq dataset^54^ containing 51,075 human and 46,667 common marmoset (*Callithrix jacchus*) cells of 30-day-old COs was re-analyzed using the processed Seurat object (DOI: 10.5281/zenodo.18480599) and core Seurat plotting functions (v5.3.0)^105^. Trajectory inference was performed using Monocle3 (v1.4.26)^106^. For aRG1–4 clusters, pseudobulk differential gene expression analysis was performed as in our previous study (Tynianskaia et al., 2026) by aggregating expression per biological replicate and cluster using Seurat’s AggregateExpression function (v5.0.3)^107^. Differential expression was tested using Seurat’s FindMarkers function with the MAST hurdle model^108^, and P-values were adjusted using the Benjamini–Hochberg method. Median pseudotime of ZNF90-positive, OVOL2-positive, and ZNF90&OVOL2-positive cells was compared using the Wilcoxon rank-sum test with continuity correction. To determine whether, and in which human clusters, co-expression of *ZNF90* and *OVOL2* was higher than expected by chance, Fisher’s exact test with Benjamini–Hochberg FDR correction was performed on each 2 × 2 co-expression contingency table. Exact adjusted P-values and sample sizes are reported in the corresponding figure legends.

